# Quantitative mass spectrometry to interrogate proteomic heterogeneity in metastatic lung adenocarcinoma and validate a novel somatic mutation CDK12-G879V

**DOI:** 10.1101/398313

**Authors:** Xu Zhang, Khoa Dang Nguyen, Paul Rudnick, Nitin Roper, Emily Kawaler, Tapan K. Maity, Shivangi Asasthi, Shaojian Gao, Romi Biswas, Abhilash Venugopalan, Constance M. Cultraro, David Fenyö, Udayan Guha

**Affiliations:** Thoracic and Gastrointestinal Oncology Branch, CCR, NCI, NIH, Bethesda, MD; Spectragen Informatics LLC, Bainbridge Island, WA; Institute for Systems Genetics and Department of Biochemistry and Molecular Pharmacology, NYU School of Medicine, New York, NY

## Abstract

Lung cancer is the leading cause of cancer death both in men and women. Tumor heterogeneity is an impediment to targeted treatment of all cancers, including lung cancer. Here, we sought to characterize changes in tumor proteome and phosphoproteome by longitudinal, prospective collection of tumor tissue of an exceptional responder lung adenocarcinoma patient who survived with metastatic lung adenocarcinoma for more than seven years with HER2-directed therapy in combination with chemotherapy. We employed “Super-SILAC” and TMT labeling strategies to quantify the proteome and phosphoproteome of a lung metastatic site and ten different metastatic progressive lymph nodes collected across a span of seven years, including five lymph nodes procured at autopsy. We identified specific signaling networks enriched in lung compared to the lymph node metastatic sites. We correlated the changes in protein abundance with changes in copy number alteration (CNA) and transcript expression. To further interrogate the mass spectrometry data, patient-specific database was built incorporating all the somatic variants identified by whole genome sequencing (WGS) of genomic DNA from the lung, one lymph node metastatic site and blood. An extensive validation pipeline was built for confirmation of variant peptides. We validated 360 spectra corresponding to 55 germline and 6 somatic variant peptides. Targeted MRM assays demonstrated expression of two novel variant somatic peptides, CDK12-G879V and FASN-R1439Q, with expression in lung and lymph node metastatic sites, respectively. CDK12 G879V mutation likely results in a nonfunctional CDK12 kinase and chemotherapy susceptibility in lung metastatic sites. Knockdown of CDK12 in lung adenocarcinoma cells results in increased chemotherapy sensitivity, explaining the complete resolution of the lung metastatic sites in this patient.

## Introduction

Lung cancer is the leading cause of cancer mortality in men and women. The identification of several actionable targets in lung adenocarcinoma, the commonest histology of non-small cell lung cancer (NSCLC), has resulted in development of targeted treatments against those targets. Although patients harboring those somatic mutational targets, such as the Epidermal growth factor receptor (EGFR) and EML4-ALK translocation respond well to the targeted agents, they eventually develop resistance. A key determinant of this acquired resistance is tumor evolution and generation of intra-tumor and inter-metastatic heterogeneity (1-3). The degree of mutational heterogeneity is highly variable within primary tumors and between primary and metastatic or recurrence sites. In some tumor types, such as recurrent glioma, the emergence of heterogeneity may be therapy related (<u>Johnson et al 2014</u>). In certain cancer types that are driven by environmental mutagens such as ultraviolet light in melanoma and smoking in lung cancer, the extent of homogenous mutational burden is greater than other caner types (4). However, there is a subgroup of lung cancer patients that demonstrate extreme intra-tumor and inter-metastatic genomic mutational heterogeneity (2, 3, 5). We have demonstrated unprecedented genomic heterogeneity in an exceptional responder lung adenocarcinoma patient who survived with metastatic lung adenocarcinoma for more than 7 years while on treatment with combination HER2-directed targeted treatment and chemotherapy. There was less than 1% similarity of somatic alterations between the lung and lymph node metastatic sites in this patient (5). Multidimensional genomic analyses of the next-generation sequencing data including, for single nucleotide variants (SNVs), insertion-deletions (in-dels), copy number alterations (CNA) and expressed variants from RNAseq can give a comprehensive view of the genomic landscape and its contribution to tumor heterogeneity. However, how this genomic heterogeneity influences the proteome and phosphoproteome is an important question that can be addressed by global mass spectrometry-based proteomics analyses. Recently, the NCI Clinical Proteomic Tumor Analysis Consortium (CPTAC) has employed mass spectrometry to analyze the proteome and phosphoproteome of colon, serous ovarian and breast primary tumors that underwent extensive genomic characterization (6-8). However understanding of temporal and spatial proteogenomic tumor evolution in metastatic lung adenocarcinoma is lacking.

Here, we have used quantitative mass spectrometry strategies to identify and quantify the proteome and phosphoproteome of sequentially acquired lung and lymph node metastatic sites across a span of 7 years during which the patient was treated with combination HER-2 directed therapy. The metastatic sites were removed by surgical excision at the time of site-specific progression of disease and the final acquisition was at the time of autopsy. Interrogation of patient-specific databases built using the whole genome sequencing data from the lung and lymph node metastatic sites identified key somatic variants at the peptide-level. We further validated the novel CDK12G879V mutation by functional analyses in lung adenocarcinoma cells and provide evidence that the presence of this mutation only in the lung, and not in lymph nodes might have resulted in sensitivity of the lung metastatic sites to chemotherapy and a “cure” of the lung metastatic sites with combination HER-2 directed targeted and standard chemotherapy in this patient.

## Methods

### 1. Tumor tissue collection by serial biopsies and at autopsy

Index patient was treated with standard of care treatment regimens or specific clinical protocols at the NIH Clinical Center or Walter Reed National Military Medical Center over a period of more than 7 years. At specific times when there was progression, patient underwent excisional biopsy of progressive lymph nodes. Patient also underwent a surgery to remove a progressive lesion in the lung (see time line, Fig 1A). Tumor tissue was frozen immediately in liquid nitrogen before transporting to the laboratory for storage at -80^°^ C. About 10-15 mg of tumor tissue was cut and lysed in 400µl of urea lysis buffer using a tissue lyser (Qiagen). Lysates were centrifuged at 14,000 rpm at 4^°^ C for 10 mins and the clear supernatants were transferred to new tubes. Protein concentrations were determined by the Modified Lowry method (BioRad).

**Figure 1.**
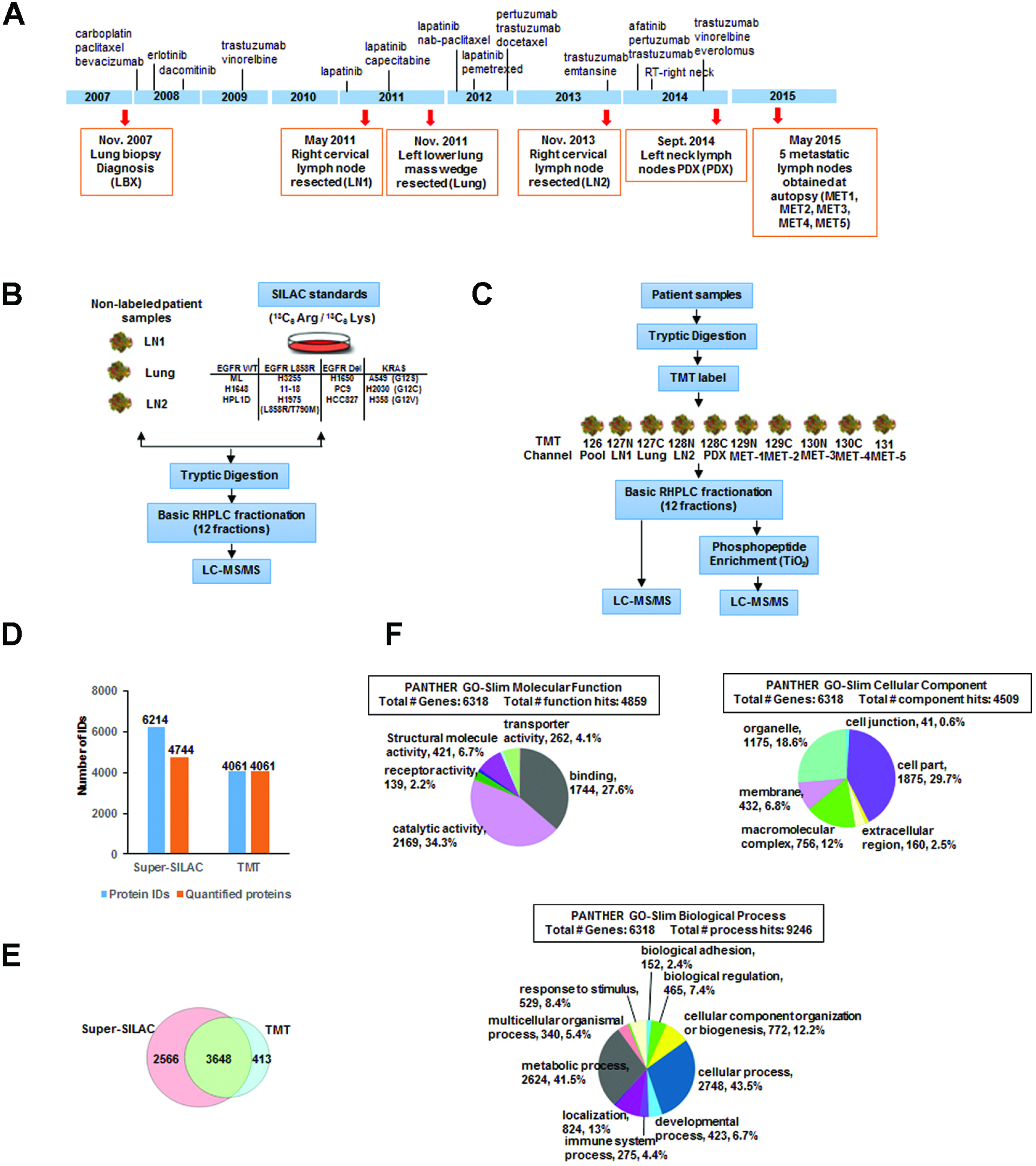
Treatment and tumor acquisition timeline, experimental workflow and summary of MS analyses. (A) The timeline of treatment history is indicated by the vertical lines along the blue bar with names of the drug used at each approximate time point. Tumor biopsies were taken at each time point (red arrow) with the date, sample description, and sample names in parenthesis (orange box). (B) Experimental workflow of super-SILAC based quantitative mass spectrometry. 12 heavy labeled (^13^C_6_ Arg, ^13^C_6_ Lys) lung adenocarcinoma cell lines were mixed at equal amount and used as standard. Then the three tumor biopsies (LN1, Lung, and LN2) were combined with the standard at equal amount, followed by tryptic digestion, high pH-RPLC fractionation, and then subjected to LC-MS/MS analysis on an Orbitrap Elite. (C) Experimental workflow of TMT based quantitative mass spectrometry. Three tumor biopsies (LN1, Lung, and LN2), one PDX sample, and five tumor autopsies from the patient were used in this experiment. All the samples were pooled at equal amount to make the reference. The digested peptides or TiO_2_ enriched phosphopeptides were TMT labelled and then combined at equal amount, followed by high pH-RPLC fractionation and then LC-MS/MS analysis on an Orbitrap Elite. (D) Bar graph showing the total number of proteins identified and quantified in both super-SILAC experiment and TMT experiment. 24% of proteins identified in super-SILAC experiment is not quantified. (E)Venn diagram showing the results of protein identification from both super-SILAC and TMT experiments. 3648 proteins were common between the two methods. (F) Pie chart represents of the distribution of identified proteins from both TMT and super-SILAC experiments together according to their molecular functions, biological processes and cellular component. Categorizations were based on information provided using the online resource PANTHER classification system.

### 2. Cell Culture and SILAC Labeling

Human lung adenocarcinoma cells were obtained from ATCC. All cells were cultured in RPMI 1640 supplemented with 10% dialyzed FBS and Penicillin/Streptomycin at 37 °C and 5% CO2. The heavy amino acids-labeled super-SILAC (*s*table *i*sotope *l*abeling with *a*mino acids in *c*ell culture) mix constituted of 2 immortalized lung epithelial cell lines (HBEC3KT and HPL1D), 1 lung adenocarcinoma cell line with wild type EGFR and KRAS (H1648), 2 EGFR^L858R^ mutant cell lines (H3255, 11-18), 1 EGFR^L858R/T790M^ cell line (H1975), 3 EGFR^DEL^ mutant cell lines (H1650, PC9, and HCC827), and 3 KRAS mutant cell lines (A549, H2030, and H358). These cell lines were labeled by culturing in RPMI with the natural lysine and arginine replaced by heavy isotope labeled amino acids, L-^13^C_6_-arginine (R6; ^13^C_6_ 98%) and L-^13^C_6_-lysine (K6; ^13^C_6_ 98%). Labeled amino acids were purchased from Cambridge Isotope Laboratories (Andover, MA). Cells were cultured for approximately seven passages in the SILAC medium for complete incorporation of the heavy isotopes. Labeling efficiency was measured by mass spectrometry analysis of tryptic peptides processed from lysates obtained from individual cell lines after 5-7 generations of growth.

### 3. Protein Extraction

Cells were lysed with urea lysis buffer (20 mM HEPES pH 8.0, 8 M urea, 1 mM sodium orthovanadate, 2.5 mM sodium pyrophosphate and 1 mM ß-glycerophosphate). Protein concentrations were determined by the Modified Lowry method (BioRad). Equal amounts of protein from lysates of each heavy labelled cell line were mixed together to constitute pooled lysate and used as a reference for the super-SILAC experiments.

### 4. Enzymatic Digestion

The protein lysate was reduced with 45 mM dithriothreitol (Sigma Aldrich, MO), alkylated with 100 mM iodoacetamide (Sigma Aldrich, MO), and subsequently digested with modified sequencing grade Trypsin (Promega, Madison, WI) at 30°C overnight. The digest was then acidified using 0.1% TFA and the peptides were desalted using solid phase extraction C18 columns (Supelco, Bellefonte, PA), and vacuum dried in a centrifugal evaporator.

### 5. TMT-Labeling

TMT10-plex amine reactive reagents (0.8 mg per vial) (Thermo Fisher Scientific) were resuspended in 41 µL of anhydrous acetonitrile (ACN), added to 200 ng protein lysates from each sample and mixed briefly on a vortexer. Reactions were proceeded at room temperature for 1 h, quenched by the addition of 8 µL of 5% hydroxylamine for 15 min and then tagged lysates were combined at equal amounts. All tumor tissues were pooled together to make the reference channel and labeled with TMT^10^-126.

### 6. Basic reversed phase liquid chromatography (RPLC) fractionation

Basic RPLC separation was performed with a XBridge C18, 100 x 2.1 mm analytical column containing 5µm particles and equipped with a 10 x 2.1 mm guard column (Waters, Milford, MA) with a flow rate of 0.25 mL/min. The solvent consisted of 10 mM triethylammonium bicarbonate (TEABC) as mobile phase A, and 10 mM TEABC in ACN as mobile phase B. Sample separation was accomplished using the following linear gradient: from 0 to 1% B in 5min, from 1 to 10% B in 5min, from 10 to 35% B in 30min, and from 35 to 100% B in 5min, and held at 100% B for an additional 3min. A total of 96 fractions were collected during the LC separation in a 96-well plate in the presence of 12.5 µL of 1% formic acid. The collected fractions were concatenated into 12 fractions and dried in a vacuum centrifuge. One tenth of the peptides was injected directly for LC-MS/MS analysis.

### 7. Titanium Dioxide (TiO_2_) Enrichment

Nine tenth of the dried peptides was dissolved in solution A containing 80% acetonitrile, 0.4% trifluoracetic acid and 3% Lactic acid and enriched with the TiO_2_ phosphopeptide enrichment spin column (Thermo Scientific). After binding, TiO_2_ spin columns were washed with solution A and thrice with 80% acetonitrile containing 0.4% trifluoracetic acid. TiO_2_ bound peptides were eluted using 5% NH_4_OH and 5% pyrrolidine and immediately acidified using trifluoracetic acid. The peptides were vacuum dried, cleaned on C18 stage-tips before LC-MS/MS analysis.

### 8. LC-MS/MS analyses

Peptides separated/fractionated by basic reversed-phase chromatography followed by TiO_2_ enrichment were analyzed on an LTQ-Orbitrap Elite interfaced with an Ultimate^™^ 3000 RSLCnano System (Thermo Scientific, San Jose, CA). The dried peptides and the enriched phosphopeptides were loaded onto a nano-trap column (Acclaim PepMap100 Nano Trap Column, C18, 5 µm, 100 Å, 100 µm i.d. x 2 cm) and separated on an Easy-spray^™^ C18 LC column (Acclaim PepMap100, C18, 2 μm, 100 Å, 75 μm i.d. × 25 cm). Mobile phases A and B consisted of 0.1% formic acid in water and 0.1% formic acid in 90% ACN, respectively. Peptides were eluted from the column at 300 nL/min using the following linear gradient: from 4 to 35% B in 60min, from 35 to 45% B in 5min, from 45 to 90% B in 5min, and held at 90% B for an additional 5min. The heated capillary temperature and spray voltage were 275°C and 2kV, respectively. Full spectra were collected from m/z 350 to 1800 in the Orbitrap analyzer at a resolution of 120,000, followed by data-dependent HCD MS/MS scans of the fifteen most abundant ions at a resolution of 30,000, using 40% collision energy and dynamic exclusion time of 30s.

### 9. MRM experiment

Heavy labeled peptides with a C-terminal ^15^N- and ^13^C-labeled arginine (^13^C_6_, ^15^N_4_) or lysine (^13^C_6_, 15N_2_) residue were purchased from Thermo Fisher Scientific. The synthetic peptides were re-constituted in 0.1% formic acid and were analyzed on the nano-chip-LC using a 1260 Infinity Series HPLC-Chip cube interface (Agilent, Palo Alto, CA) coupled to a 6495-triple quadrupole mass spectrometer (Agilent). A large capacity chip system (G4240-62010) consisting of a 160 nl enrichment column and a 150 mm*75 um analytical column (Zorbax 300SB-C18, 5 um, 30 A pore size) was used. Mobile phase A consisted of 95% water and 0.1% FA, and mobile phase B consisted of 95% acetonitrile in 0.1% FA. A flow rate of 3ul/min was applied for sample loading by the capillary pump and 600 nl/min for the analytical separation through the nano-pump. A 25-minute gradient (0 min, 0% B; 1 min, 5% B; 17 min, 40% B; 20 min, 70% B; 21 min, 80% B; 25 min 0% B) was used for the chromatographic separation of the target peptides. A spray voltage of 1850 V was applied and quadrupoles 1 and 3 were run at 0.7 FWHM resolution. Individual injections were carried out to select optimum precursors and intense product ions for each synthetic peptide. The top transitions were selected based on the presence of intense y-ions at m/z greater than the precursor. In case high m/z y-ions were not seen, other more abundant ions were chosen. All the raw data were imported in Skyline 3.7 and manually reviewed to delete any poorly performing transitions. Each peptide consisted of two to five transitions. The optimization of collision energies was done by using the values calculated by Skyline for the monoisotopic precursor and product masses for the Agilent 6495 system. The best performing transitions were combined in one scheduled MRM method with a 25-min gradient and a 2-minute retention time window, using retention times extracted during the method refinement stage. For endogenous target peptide identifications, heavy labeled synthesized peptides were spike in as an internal standard to different tissue samples. All the raw data files were imported in Skyline 3.7 and data annotations were manually inspected. Peak Area Ratios (PAR) of light endogenous signals to the heavy internal standards were exported to MS excel for further analysis of mean, standard deviation and % co-efficient of variation.

### 10. Tumor Specific Database Construction

Customized patient specific database was created by QUILTS (9) (quilts.fenyolab.org) using RefSeq as the reference protein sequence database. (i) a BED file containing RNA-Seq predicted junctions; VCF files containing (ii) somatic variants and (iii) germline variants; and (iv) a fusion file containing all predicted fusion genes were used as inputs. QUILTS creates tumor-specific databases by enumerating all possible protein sequence variation that results from the genomic and transcriptomic sequence variation including single amino acid variants, introduced and removed stop codons, alternative splicing, fusion proteins, expression of unannotated parts of the genome. Our database for this study contained a total of 71,503 entries.

### 11. Data Analysis

Peptides and proteins were identified and quantified using the Maxquant software package (version 1.5.3.30) with the Andromeda search engine (10, 11). MS/MS spectra were searched against the customized patient specific database and quantification was performed using default parameters for 3s-SILAC or TMT10plex in MaxQuant. The parameters used for data analysis include trypsin as a protease with two missed cleavage sites allowed. Carbamidomethyl cysteine was specified as a fixed modification. Phosphorylation at serine, threonine and tyrosine, deamidation of asparagine and glutamine, oxidation of methionine and protein N-terminal acetylation were specified as variable modifications. The precursor mass tolerance was set to 7 ppm and fragment mass tolerance to 20ppm. False discovery rate was calculated using a decoy database and a 1% cut-off was applied to both peptide table and phosphosite table.

Normalized ratios from super-SILAC experiment or corrected intensities of the reporter ions from TMT labels were obtained from the MaxQuant search. For the TMT experiment, relative ratios of each channel to the reference channel were calculated. Perseus (version 1.5.5.3) was used to view and further analyse the data. Hierarchical clustering of proteins and phosphosites were obtained in Perseus using log ratios or log intensities of protein and phosphorylation abundance. The Protein-Protein Interaction (PPI) maps of the phosphosite clusters were imported from the “STRING: protein query” module of the cystoscope software (San Diego, CA, USA, version 3.4.0) (12) with the confidence cutoff of 0.80. These maps were analyzed for functional enrichment of the gene ontology biological process categories using the ClueGO 2.2.6 plugin (13)with the kappa statistic >=0.4, a two-sided hypergeometric test for enrichment with Bonferroni step down method for correction of the multiple hypothesis testing. A p-value of 0.001 was used as the cut-off criterion.

### 12. Correlation between mRNA expression and protein abundance

We correlated mRNA expression and protein abundance within each of two metastatic tumor sites (L and LN1) and the PDX. We used RNA-seq FPKM values and protein intensity measurements from TMT and SILAC experiments to estimate mRNA and protein abundance, respectively. For each tumor site, we calculated the Spearman correlation coefficient between log2 values of FPKM and protein intensity values for genes that had measurements in both mRNA and protein (number of genes for TMT experiments were: L, 3853; LN1, 3880; PDX, 3791, and number of genes for SILAC measurements were: L, 5890; LN1, 5920).

### 13. Copy number variation analysis

We used OncoScan CNV FFPE Assay Kit (Affymetrix) to perform copy number variation (CNV) analysis. Copy number data was processed and normalized using OSCHP-TuScan algorithm, which determines allele specific copy number and estimates ploidy. Copy number log2 ratios of each tumor site was estimated with Nexus Express for OncoScan. Regions of copy number gains or losses across all metastatic tumor sites were matched based on the location of genes in each region. A CNV heatmap was generated based on hierarchical clustering analysis using Euclidean distance.

### 14. Mutant peptide validation

Confirmatory analysis of MS2 spectra was performed separately and following MaxQuant analysis using mutant-specific database. First, Oribtrap Elite HCD spectra were converted to MGF format using the NIST program ReAdW4Mascot2.exe (v20130604a) (http://chemdata.nist.gov/dokuwiki/doku.php?id=peptidew:pepsoftware). Parameters for conversions were as follows: -c -ChargeMgfOrbi -FixPepmass -MaxPI -MonoisoMgfOrbi - NoPeaks1 -PIvsRT -sep1 -TolPPM 20 -NoMzXml. These settings include the -FixPepmass option which reassess the monoisotopic peak, assigned by XCalibur^™^, using the previous and next MS1 scans, changing the PEPMASS value if a better assignment can be made.

All fragmentation spectra from the Super SILAC experiments were searched using Mascot 2.5 (Matrix Science, Ltd., London, UK) and MS-GF+ (v20140716) against the patient-specific database created using the QUILTS pipeline. Mascot searches were run using a 20ppm precursor mass tolerance (monoisotopic) and a 0.05 Da fragment mass tolerance, allowing for up to 2 missed cleavages and considering only fully tryptic peptides. Quantitation mode was set to SILAC K+6; R+6. Carbamidomethylation of C was set as a fixed modification and oxidized methionine as variable. MS-GF+ settings were the following: 10 ppm precursor mass tolerance, -m 3, -inst 1, - ntt 2, -tda 1, -ti 0, 0 -max Length 60 -min Length 7. These settings were specific for Orbitrap HCD spectra, performed target-decoy-analysis, did not allow for isotopic matching, and excluded peptides longer than 60 and shorter than 7 amino acids long. Carbamodomethyl of C was used as a fixed modification and heavy versions of K and R (+6.020), oxidation of M, and protein N-terminal acetylation were used as variable modifications. MZID files were converted to TSV using the MzIDToTSV function of the program.

Next, spectra from identifications of variant peptides identified by MaxQuant were extracted following removal of peptides that did not contain variant sequences (e.g., those giving rise to a K or R adjacent to the start of the peptide sequence). In total, 833 spectra corresponding to 105 sequences (of the 198 identified by MaxQuant) were analyzed. All spectra were required to have corresponding identifications better than the Mascot Identity Threshold (95%) or a Q Value < 0.01 (MS-GF+) per file. We found no disagreements between Mascot and MS-GF+ for these spectra, and when both search engines agreed, the Mascot identification information was kept.

Since variant sequences, in particularly those that are unknown (i.e., not found in dbSNP) or somatic are rare, additional steps were taken to ensure the quality of the peptide-spectrum matches. These steps were performed according to (9) and in consideration of Nesvishskii et al. (14). First, all fragmentation spectra were labelled and the fraction of matched MS2 intensity was calculated for each. These values provide a good measure of the purity of a spectrum, and there is a correlation with a low unmatched intensity and correct assignments. Additionally, sequencing “gaps” were analysed (9). A gap is defined as missing fragmentation evidence between adjacent residues. This analysis considers b- and y-type ions only (including neutral losses of water and ammonia) and is based on the observation that large gaps such as the complete absence of one or the other ion series can give rise to good matching identifications but often correspond to incorrect matches and cases where the correct peptide sequence is not in the target database. For our analysis, we removed all spectra with unmatched abundance of >50% and those with any sequence gap >2 (9).

In addition to Mascot and MS-GF+ searches, X! Tandem searches of the 833 putative variant peptide spectra were run. These searches were run to make use of X! Tandem’s refinement mode options as well as search the common contaminants database cRAP. In particular, X!Tandem can search for single amino acid substitutions as well as Single Amino acid Polymorphism (SAPs) indexed from dbSNP. Settings for X! Tandem search were the following: version-Vengance (2015.12.15.2), 20 ppm precursor tolerance, 20 ppm fragment mass tolerance, fixed cabamidomethyl C, variable oxidized M and deamidated N. For refinement searching, carbamylation of K and/or the peptide N-terminus was optionally allowed, as was dioxidation of M. Substitution of up to 1 amino acid and SAP searching were also turned on. Agreement between the X! Tandem analysis and the Mascot/MS-GF+ analysis was very good. However, X! Tandem did not confidently identify 51 of the 833 spectra, returning no match. Additionally, we found peptides where X! Tandem disagreed. One case, preferring a deamidated N rather than the N>D amino acid change. Because of this result all matches to N->D peptides were excluded from further analysis. Additionally, for another peptide, the top-ranking X! Tandem match was an amino acid change to an alternate reference peptide over the variant present in the database. Since the variant was found by genomic sequencing, the variant identification was kept over the substitution.

### 15. Targeted analysis using multiple reaction monitoring (MRM)

Heavy labeled peptides synthesized with a C-terminal ^13^C_6_^15^N_4_-labeled arginine or ^13^C_6_ ^15^N_2_ -labeled lysine were purchased from Thermo Fisher Scientific. The synthetic peptides were re-constituted in 0.1% formic acid and were analyzed on the nano-chip-LC using a 1260 Infinity Series HPLC-Chip cube interface (Agilent, Palo Alto, CA) coupled to a 6495-triple quadrupole mass spectrometer (Agilent). A large capacity chip system (G4240-62010) consisting of a 160 nl enrichment column and a 150 mm*75 µm analytical column (Zorbax 300SB-C18, 5 µm, 30 A pore size) was used. Mobile phase A consisted of 95% water and 0.1% FA, and mobile phase B consisted of 95% acetonitrile in 0.1% FA. A flow rate of 3µl/min was applied for sample loading by the capillary pump and 600 nl/min for the analytical separation through the nano-pump. A 25-minute gradient (0 min, 0% B; 1 min, 5% B; 17 min, 40% B; 20 min, 70% B; 21 min, 80% B; 25 min 0% B) was used for the chromatographic separation of the target peptides. A spray voltage of 1850 V was applied and quadrupoles 1 and 3 were run at 0.7 FWHM resolution. Individual injections were carried out to select optimum precursors and intense product ions for each synthetic peptide. The top transitions were selected based on the presence of intense y-ions at m/z greater than the precursor. In case high m/z y-ions were not seen, other more abundant ions were chosen. All the raw data were imported in Skyline 3.7 and manually reviewed to delete any poorly performing transitions. Each peptide consisted of four or five transitions. The optimization of collision energies was done by using the values calculated by Skyline for the monoisotopic precursor and product masses for the Agilent 6495 system. The best performing transitions were combined in one scheduled MRM method with a 25-min gradient and a 2-minute retention time window, using retention times extracted during the method refinement stage. For endogenous target peptide quantification, heavy labeled synthesized peptides were spiked-in as an internal standard to the tryptic peptides of different tissue samples. All the raw data files were imported in Skyline 3.7 and data annotations were manually inspected. Peak Area Ratios (PAR) of light endogenous signals to the heavy internal standards were exported to MS excel for further analysis of mean, standard deviation and % co-efficient of variation.

### 16. CRISPR-Cas9-mediated knockout of CDK12 and cell growth analysis

A549 cells were infected with lentivirus expressing Cas9. Clones were obtained after antibiotic selection and one of the clones with higher expression of Cas9 was selected for experiments based on Western blot detection of Cas9. A549-Cas9 cells were then transduced with lentivuruses expressing CRISPR sgRNAs that target exon 1 and exon 2 of *CDK12*. After puromycin selection, clones were picked and expanded. Cells were lysed with 1x modified RIPA buffer, and immunoblotted for CDK12 expression. Two clones, A549-Cas9-261-7 and A549-Cas9-264-5, in which the sgRNAs target exon 1 and exon 2 of *CDK12*, respectively, were selected for further functional analysis based on the knockdown of CDK12 expression.

For chemotherapy treatment and survival analysis, cells were trypsinized with 0.25% trypsin in EDTA, and spun down at 1,000 rpm for 5 mins at room temperature. Cells were re-suspended in RPMI medium with 10% FBS, 1% pen/strep, counted with trypan blue exclusion and plated 4,000 cells per well in a 96 well-plate. Cells were allowed to settle overnight before drug treatment. 10x stock of camptothecin (Cell Signaling Technology) was prepared fresh for each experiment. 10uL of camptothecin was added to 90uL of RPMI media already present in each well. Cells were incubated at 37°C, 5% CO2 for 48 or 72 hours. After 48 or 72 hours of treatment, all media was removed, and 50uL of 1x Promega CellTiter-Glo^®^ Luminescent Cell Viability Assay reagent was added to each well. Luminescence was measured with a SpectraMax M5 microplate reader and recorded by SoftMax Pro 5.4.1. Raw luminescence was normalized and plotted with MS Excel.

## Results

### Brief clinical history, tumor tissue procurement and summary of protein identification and quantification

A 50-year old African American never smoker was diagnosed with metastatic lung adenocarcinoma with metastases to lung and lymph nodes. Patient was treated with combination chemotherapy followed by HER2-directed therapy either alone or in combination with various chemotherapy regimens over 7 years. Two case reports (15, 16) and a comprehensive genomics analysis of multiple tumor biopsies across his 7-year treatment course have been reported demonstrating extreme heterogeneity between lung and lymph node metastatic sites in this patient (5). Here, we sought to determine the temporal and spatial heterogeneity of the proteome and report a comprehensive quantitative proteomics analysis on different tumors metastatic to lymph nodes (LN1, LN2 and PDX) obtained by sequential excisional lymph node biopsies, 5 distinct lymph node metastases obtained at autopsy (MET1-5), and a progressive metastatic lung tumor obtained by wedge resection surgery (L) (**Fig. 1A**). The patient-derived xenograft (PDX) was generated from one of the lymph node biopsies from the left neck. The patient was treated with combination chemotherapy followed by targeted therapy in the form of HER2-directed treatments with/without chemotherapy throughout his course of more than 7 years as depicted in the timeline (**Fig. 1A**). Both super-SILAC and TMT10plex-based quantitative proteomics strategies were employed to explore the tumor-specific and temporal heterogeneity of the expressed proteomes. In the super-SILAC strategy, twelve “heavy” amino acids-labelled lung adenocarcinoma and immortalized lung epithelial cell lines were pooled at equal amounts and used as the standard for “spike-in”. Lysates from three distinct metastatic tumors (LN1, Lung and LN2) were combined with the “heavy” standard lysate at equal amounts for further sample processing and mass spectrometry analysis (**Fig. 1B**). The TMT-based quantitative proteomics strategy was used for the quantification of the proteome and phosphoproteome of nine distinct tumor metastases, including two lymph nodes (LN1 and LN2), one lung tumor (L), one PDX derived from a lymph node metastasis, and five lymph node metastases (MET1-5) procured at autopsy (**Fig. 1C**). Lysates from the nine samples were pooled at equal amounts and labelled with TMT^126^ to be used as the reference. After trypsin digestion and basic reverse-phase HPLC fractionation, tandem mass spectrometry data at high accuracy and resolution was acquired on an Orbitrap Elite mass spectrometer. A total of 6214 and 4061 proteins were identified from super-SILAC and TMT experiments, respectively (**Fig. 1D**). More than 4000 proteins were quantified from both experiments. 3648 proteins were identified and quantified in both experiments (**Fig 1E)**. PANTHER analysis gives the protein distribution based on their molecular functions, biological processes and cellular components. More than 2000 proteins had catalytic activity that include kinases, phosphatases and metabolic enzymes (**Fig. 1F**).

### Correlation and distribution of protein abundance ratios and clustering analysis reveals more similarity of protein abundances between the lung and LN2 metastases

Next, we compared the protein quantification results between the super-SILAC approach and the TMT approach. Correlations between the two different quantification approaches and three different samples were plotted. The protein ratios across three different samples (LN1, L and LN2) were well correlated between super-SILAC and TMT approaches (**Fig. 2A**). Correlation of the protein SILAC ratios demonstrated that the protein abundance of LN2 correlated better with lung and LN1 (R^2^ = 0.68 and 0.67, respectively). However, protein abundance ratios of LN1 correlated less with that of the lung (R^2^ = 0.53), indicating that the lung tumor was more similar to the LN2 lymph node that was procured 2 years after the lung surgery than LN1 that was procured 6 months before the surgery. From the SILAC protein ratio distribution, we observed that the large majority of proteins were unchanged (**Fig. 2C**). Around 12% of proteins from each tumour biopsy sample were differentially expressed > 2-fold relative to the heavy labelled standard lysate mixture. 8-9% of the proteins were differentially expressed < 2-fold in LN2 and lung; however, in LN1, 19% of the proteins were differentially expressed < 2-fold. The number of proteins differentially expressed <2-fold in LN1 is twice that of LN2 and lung, indicating that more proteins had lower abundance in LN1 compared to LN2 and lung. Hierarchical clustering of proteins based on their “Super-SILAC” abundance ratios showed that LN2 and lung clustered together, providing further evidence of greater heterogeneity in the proteome of LN1 compared to that of Lung and LN2 (**Supplementary Fig. 1A**). We identified many extracellular proteins and proteins involved in immune effector processes with increased expression compared to the standard lysate. Some of the membrane proteins involved in protein glycosylation and GTPase activity had lower expression level in LN1 compared to LN2 and lung.

**Figure 2.**
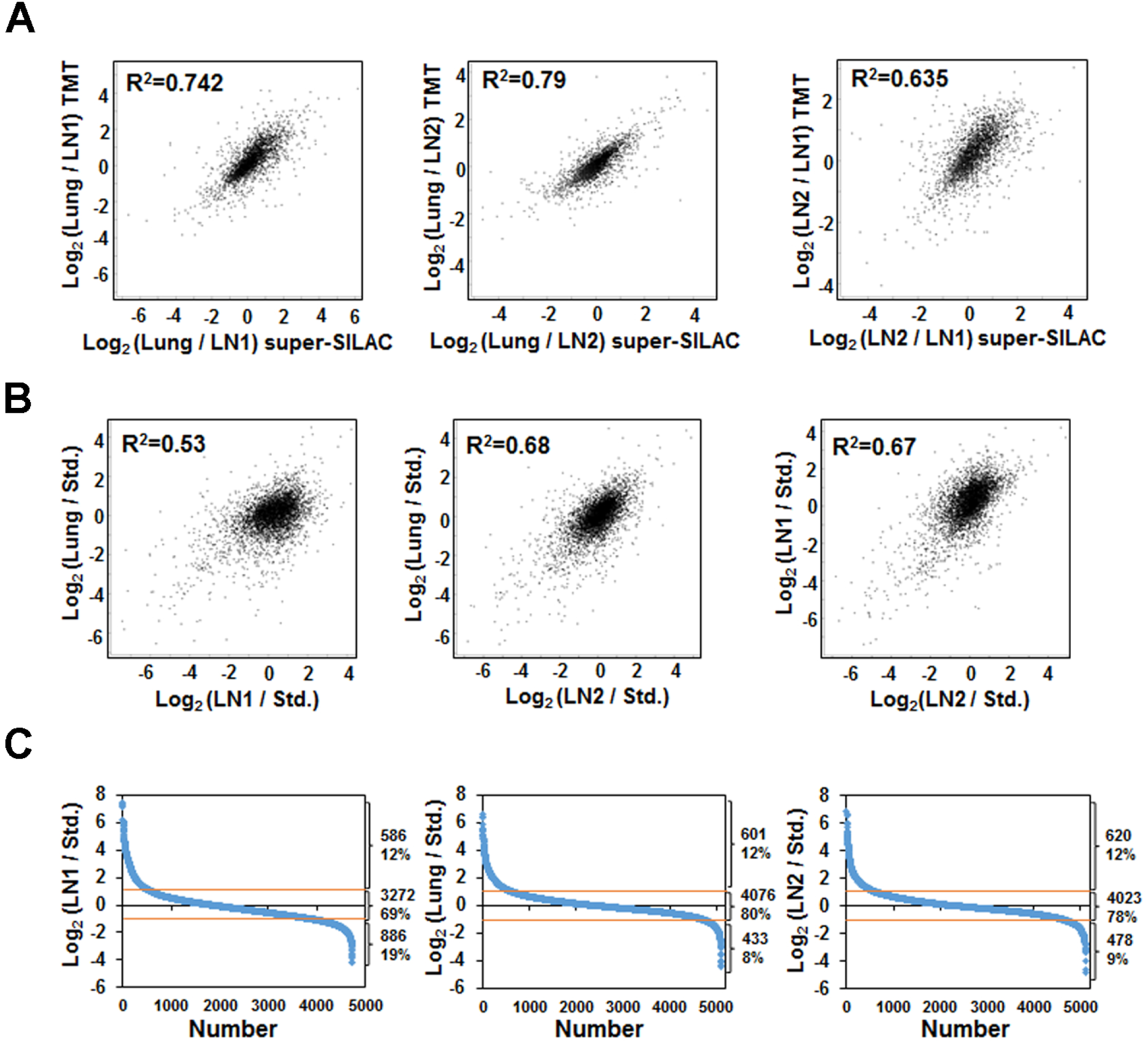
Correlation of the protein quantification. (A) Correlation between TMT and super-SILAC experiment. X-axis is the Log 2 ratio between two samples from super-SILAC approach, and y-axis is the Log 2 ratio between the same two samples from TMT approach. (B) Correlation between the three tumour biopsies samples from the results of super-SILAC experiment. It is based on the Log 2 ratio of the tumour samples and the heavy labelled super-SILAC standard. (C) Distribution of proteins quantified for the three biopsies samples (LN1, Lung, and LN2) based on the ratio against the heavy labelled standard in the super-SILAC experiment. 19% of proteins down-regulated and ∼70% of proteins remain unchanged in LN1, while 8-9% of proteins down-regulated and ∼80% proteins remain unchanged in Lung and LN2.

PCA analysis of TMT protein ratios across all metastatic sites demonstrated that lung and LN2 metastatic sites were more similar to each other and LN1. LN1, the first progressive metastatic lesion removed was different than all other metastatic sites (**Supplementary Fig. 2A**). Pair-wise correlation analysis of the TMT protein ratios showed strong correlation of specific autopsy lymph nodes, MET4 and MET5 (R^2^ value 0.845). Overall, there was poor correlation of TMT protein ratios across all metastatic sites except between lung and LN2 and between autopsy lymph nodes (**Supplementary Fig. 2B).** Hierarchical clustering of protein abundance ratios of each tumor lysate against the pooled lysate from the TMT experiment also demonstrated that LN2 and lung clustered together. LN1 protein abundance ratios clustered with three lymph nodes obtained at autopsy and was quite separate from the other two lymph node metastases (**Supplementary Fig. 1B**).

### Differentially expressed classes of proteins and enrichment of functional pathways between the lung and the two lymph node metastases, LN1 and LN2

We interrogated specific classes among the identified proteins in our “super-SILAC” quantitative mass spectrometry experiments and identified proteins encoding many cancer genes, kinases, phosphatases, transcription and translation regulators with differential expression in the lung, LN1 and LN2 metastatic sites (**Table 1**). Protein abundances of several cancer genes that were upregulated in the lung compared to the lymph node metastatic sites were ERBB2, CTNNB1, IDH2, MLH1, NDRG1, and NF1 and that were significantly downregulated in the lung compared to the lymph nodes were B2M, CDKN2A, FNBP1, MAX and SYK. Among the kinases that had increased abundance in the lung compared to the lymph nodes were CCNK, MAPK3, MAP2K3, PAK1, PDK3, and YES1 and the kinases that had decreased abundance in the lung were AURKA, CKB, STK26 and STK4. DUSP23 and PPFIA1 levels were elevated in the lung metastatic sites among phosphatases. Among the transcriptional regulators that were significantly upregulated in the lung metastatic site compared to LN1 were JUNB and SMARCD2. Interestingly, the levels of these two proteins were similar in the lung and LN2 that was excised more than two years after the removal of the lung metastasis.

**Table 1.** Selected proteins that changed 3-fold between any of the two biopsy tumor samples from super-SILAC experiment.

| Gene | Ratio Lung/LN1 | Ratio Lung/LN2 | Ratio LN2/LN1 | Gene | Ratio Lung/LN1 | Ratio Lung/LN2 | Ratio LN2/LN1 |
| --- | --- | --- | --- | --- | --- | --- | --- |
| **Cancer genes** |  |  |  | **Phosphatases** |  |  |  |
| B2M | 0.2 | 0.5 | 0.4 | BPGM | 0.3 | 0.4 | 0.7 |
| CDH1 | 3.5 | 1.6 | 2.2 | DUSP23 | 4.5 | 1.5 | 3.1 |
| CDK12 | 13.8 | 6.8 | 2.0 | NT5C | 1.2 | 4.1 | 0.3 |
| CDKN2A | 0.2 | 0.07 | 3.1 | NT5C3A | 0.3 | 0.5 | 0.6 |
| CTNNB1 | 3.1 | 2.2 | 1.4 | NUDT2 | 0.6 | 0.3 | 2.1 |
| ERBB2 | 70.3 | 21.8 | 3.2 | NUDT3 | 0.6 | 0.2 | 3.2 |
| FNBP1 | 0.2 | 0.6 | 0.3 | NUDT9 | 2.2 | 3.3 | 0.7 |
| IDH2 | 7.1 | 2.6 | 2.8 | OCRL | 0.2 | 0.6 | 0.3 |
| LCP1 | 0.2 | 0.4 | 0.5 | PPFIA1 | 3.7 | 6.4 | 0.6 |
| MAX | 0.3 | 0.5 | 0.5 | PPP1R12C | 0.2 | 0.6 | 0.4 |
| MLH1 | 3.6 | 1.8 | 2.0 | PPP1R1B | 3.4 | 2.3 | 1.5 |
| MYH11 | 0.1 | 0.6 | 0.2 | PPP1R7 | 0.3 | 0.4 | 0.6 |
| NDRG1 | 2.8 | 3.3 | 0.8 | PTPN6 | 0.1 | 0.3 | 0.3 |
| NF1 | 10.9 | 3.7 | 3.0 | TIMM50 | 0.4 | 1.3 | 0.3 |
| NFIB | 0.9 | 0.2 | 5.5 | **Transcription regulator** |  |  |  |
| SYK | 0.3 | 0.5 | 0.5 | ANKLE2 | 6.7 | 6.5 | 1.0 |
| TP53 | 0.5 | 2.4 | 0.2 | ARHGAP35 | 3.5 | 1.2 | 2.9 |
| **Kinases** |  |  |  | ASCC1 | 2.9 | 0.9 | 3.4 |
| AURKA | 0.3 | 0.3 | 0.8 | BASP1 | 0.4 | 1.6 | 0.2 |
| CCNK | 4.9 | 2.6 | 1.9 | BTAF1 | 0.6 | 2.9 | 0.2 |
| CDK2 | 3.7 | 0.8 | 4.9 | CDKN2A | 0.2 | 0.1 | 3.1 |
| CHKB | 3.4 | 2.6 | 1.3 | CGGBP1 | 0.2 | 0.6 | 0.4 |
| CKB | 0.2 | 0.2 | 1.1 | CTNNB1 | 3.1 | 2.2 | 1.4 |
| CKMT1A | 0.3 | 0.04 | 7.0 | GTF2A1 | 0.4 | 1.3 | 0.3 |
| CSNK1D | 5.1 | 2.6 | 1.9 | IFI16 | 0.3 | 0.8 | 0.3 |
| DGKA | 0.3 | 0.7 | 0.4 | IRF9 | 0.8 | 0.1 | 5.9 |
| MAP2K3 | 3.8 | 1.9 | 2.0 | JUN |  | 3.0 |  |
| MAP4K5 | 4.9 | 1.0 | 5.1 | JUNB | 9.0 | 1.3 | 7.1 |
| MAPK3 | 4.5 | 3.2 | 1.4 | KANK2 | 0.3 | 1.2 | 0.2 |
| MPP1 | 0.3 | 0.4 | 0.8 | LBH | 0.2 |  |  |
| NME3 | 5.3 | 3.4 | 1.6 | LPXN | 0.1 | 0.3 | 0.3 |
| PAK1 | 4.9 | 3.9 | 1.3 | MAX | 0.3 | 0.5 | 0.5 |
| PDK3 | 4.3 | 1.8 | 2.5 | MECP2 | 0.1 | 0.6 | 0.2 |
| PI4K2A | 3.0 | 0.1 | 25.5 | MED27 | 3.7 | 1.0 | 3.8 |
| PRKAR2B | 0.2 | 0.4 | 0.4 | NFIB | 0.9 | 0.2 | 5.5 |
| PRKCA | 0.3 | 0.6 | 0.5 | PTGES2 | 2.2 | 0.7 | 3.4 |
| SRPK2 | 0.9 | 0.2 | 4.6 | PTRF | 0.2 | 0.9 | 0.2 |
| STK26 | 0.3 | 0.5 | 0.7 | RELB | 0.2 | 0.3 | 0.6 |
| TK1 | 0.05 | 1.0 | 0.05 | SMARCD2 | 4.8 | 1.1 | 4.6 |
| YES1 | 4.4 | 2.8 | 1.6 | TEFM | 3.0 | 0.9 | 3.4 |
| STK4 | 0.1 | 0.5 | 0.3 | TFB1M |  | 4.3 |  |
| **Translation regulators** |  |  |  | THOC1 | 0.5 | 1.8 | 0.3 |
| EEF1A2 | 2.6 | 3.7 | 0.7 | TLE1 | 4.1 | 2.0 | 2.1 |
| IGF2BP2 | 0.4 | 2.2 | 0.2 | UXT | 1.6 | 0.5 | 3.5 |
| PATL1 | 0.05 | 0.1 | 0.4 | YBX3 | 0.04 | 0.1 | 0.3 |

Next, we examined the function of proteins differentially expressed greater than 2-fold between either of the two lymph node metastases (LN1 vs LN2) or between either lymph node and lung metastases in the super-SILAC experiment. Ingenuity Pathway Analysis (IPA) was performed on proteins that had a “Super-SILAC” ratio of 3-fold. Among the top activated canonical pathways represented by the proteins which had at least 3 fold change in lung compared to either of the lymph nodes were activation of IRF by cytosolic pattern recognition receptors, Rac signalling, HGF signalling, ERBB and Ephrin B signalling and among the top inhibited pathways were Interferon, Sirtuin, PKA and TP53 signalling pathways (**Fig. 3A**). The heatmaps of protein ratios of lung versus either of the lymph nodes in three selected pathways namely ERBB, Sirtuin and PKA signalling pathways showed differences in abundance of proteins in these specific pathways between the lung and lymph node sites (**Fig. 3B-D**). The expression of ERBB2, MAPK3, MAP2K2 and PAK1 in the ERBB signalling pathway was higher in lung compared to the lymph nodes.

**Figure 3.**
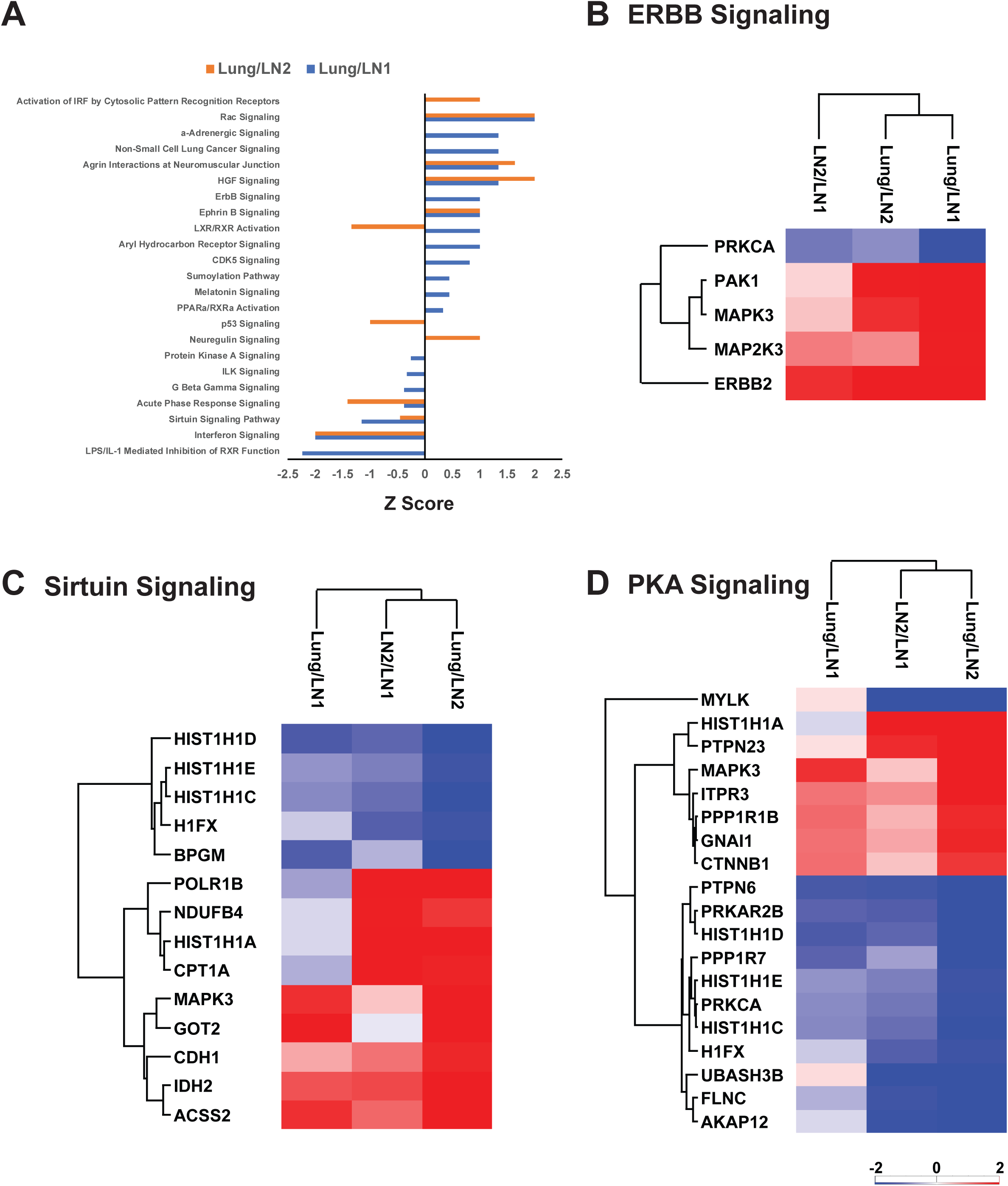
Figure 3. IPA analysis of significant changed proteins. (A) Selected top canonical pathways represented by proteins with 3-fold changes between lung and either lymph nodes. X-axis is the Z-score of the enriched pathways. Positive Z-score means the pathway is activated and negative Z-score means the pathway is inhibited. (B) Heatmap of the significant changed proteins in ERBB signalling. (C) Heatmap of the significant changed proteins in Sirtuin signalling. (D) Heatmap of the significant changed proteins in PKA signalling.

### Clustering of metastases based on copy number alteration and correlation of gene expression and protein abundance data

Hierarchical clustering of the CNA data showed that copy number of genes across many chromosomal locations from the lung tumor was quite distinct compared to rest of the lymph nodes, confirming significant metastatic site-specific copy number heterogeneity (**Fig. 4A**). Interestingly, certain lymph node metastases had more similar CNA than others, and hence clustered together. The PDX that was generated from a lymph node was more similar to LN2 than other lymph nodes. RNA was available for three samples-LN1, lung and PDX. Transcriptome sequencing (RNA-seq) was performed to quantify the transcript expression in these three metastases. Since we had the protein expression data by super-SILAC approach for LN1 and lung and by TMT approach for LN1, lung and PDX we performed Spearman’s correlation of the RNA expression values with either the super-SILAC or TMT protein abundance ratios. The correlation coefficients (R^2^) between the RNA expression values and protein abundance ratios ranged between 0.5162 for lung (L) using the super-SILAC ratio (**Fig. 4B**) and 0.281 for LN1 using the TMT protein ratio (**Fig. 4C**).

**Figure 4.**
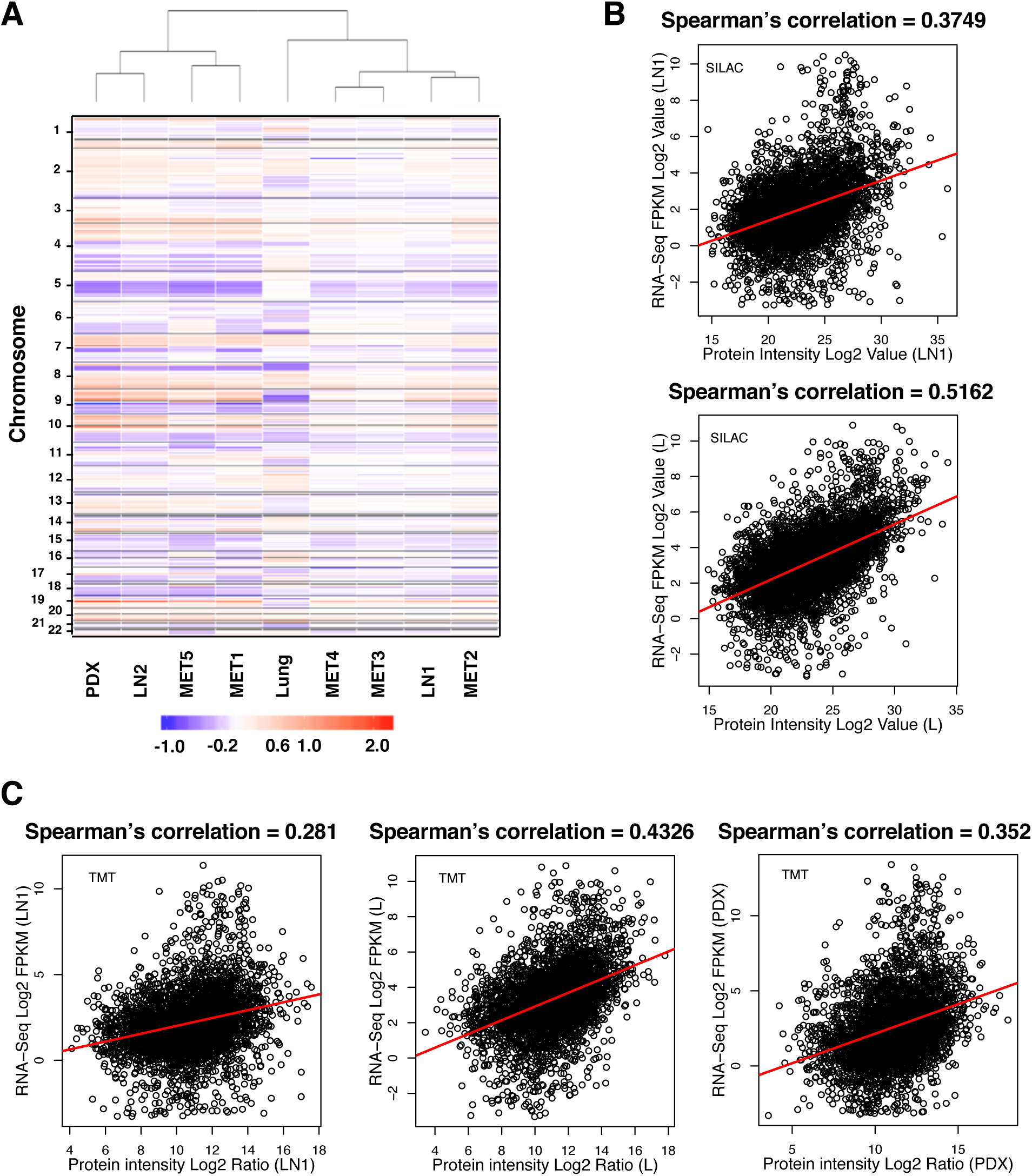
Heterogeneity in copy number alteration analysis and correlation of RNA-seq and protein estimation data. (A) Hierarchical clustering by copy number across the genome across all tumors (at cytoband resolution). Losses (purple) and gains (red) in log2 scale are depicted relative to mean ploidy. (B) Correlation of tissue LN1 and Lung between proteome results of super-SILAC and RNA-seq results. X axis is the Log 2 intensity; Y axis is the Log 2 RPKM value. (C) Correlation of tissue LN1, Lung, and PDX between proteome results of TMT and RNA-seq results. X axis is the Log 2 intensity; Y axis is the Log 2 RPKM value.

### Heterogeneity across different metastases based on global protein phosphorylation levels

Next, we examined global protein phosphorylation of TMT-labelled lysates of different metastases using TiO_2_-based enrichment strategy. The PCA plot of phosphorylation ratios shows that the autopsy lymph nodes cluster together. Similar to the protein quantitation ratios, LN2 was again closer to lung (L) compared to LN1 (**Supplementary Fig. 3A**). Pair-wise correlation analysis of the phosphorylation ratios showed strong correlation of autopsy lymph nodes (R^2^ values ranging from 0.486 to 0.835) (**Supplementary Fig. 3B**). Hierarchical clustering using the quantitative phosphorylation data across the tissue samples confirmed the similarity of the 5 autopsy lymph nodes. LN1 clustered with LN2, and the lung and the PDX lymph nodes were distinct (**Fig 5A**). There was a distinct cluster of proteins whose phosphorylation was inhibited in the autopsy lymph nodes (Cluster 662, **Fig. 5A**). Network analysis of these proteins showed that regulation of Rho protein signal transduction, cell-cell junction organization, regulation of mRNA splicing via spliceosome, spliceosome complex activity, and nuclear-transcribed mRNA catabolic process were the top networks represented by the hypo-phosphorylated proteins in autopsy samples (**Fig. 5B**). There were several proteins that had increased phosphorylation in the autopsy lymph nodes (Clusters 633, **Fig. 5A**). Enriched networks among these proteins included cellular component disassembly involved in execution phase of apoptosis, regulation of mRNA processing and mRNA export from nucleus (**Fig. 5C**). There were proteins that were hyper-phosphorylated across all the tissue samples (Cluster 616, **Fig. 5A**) and the networks that were enriched among these proteins included spliceosome complex assembly, cellular component disassembly involved in execution phase of apoptosis, establishment of spindle orientation and mRNA export from nucleus (**Fig. 5D**).

**Figure 5.**
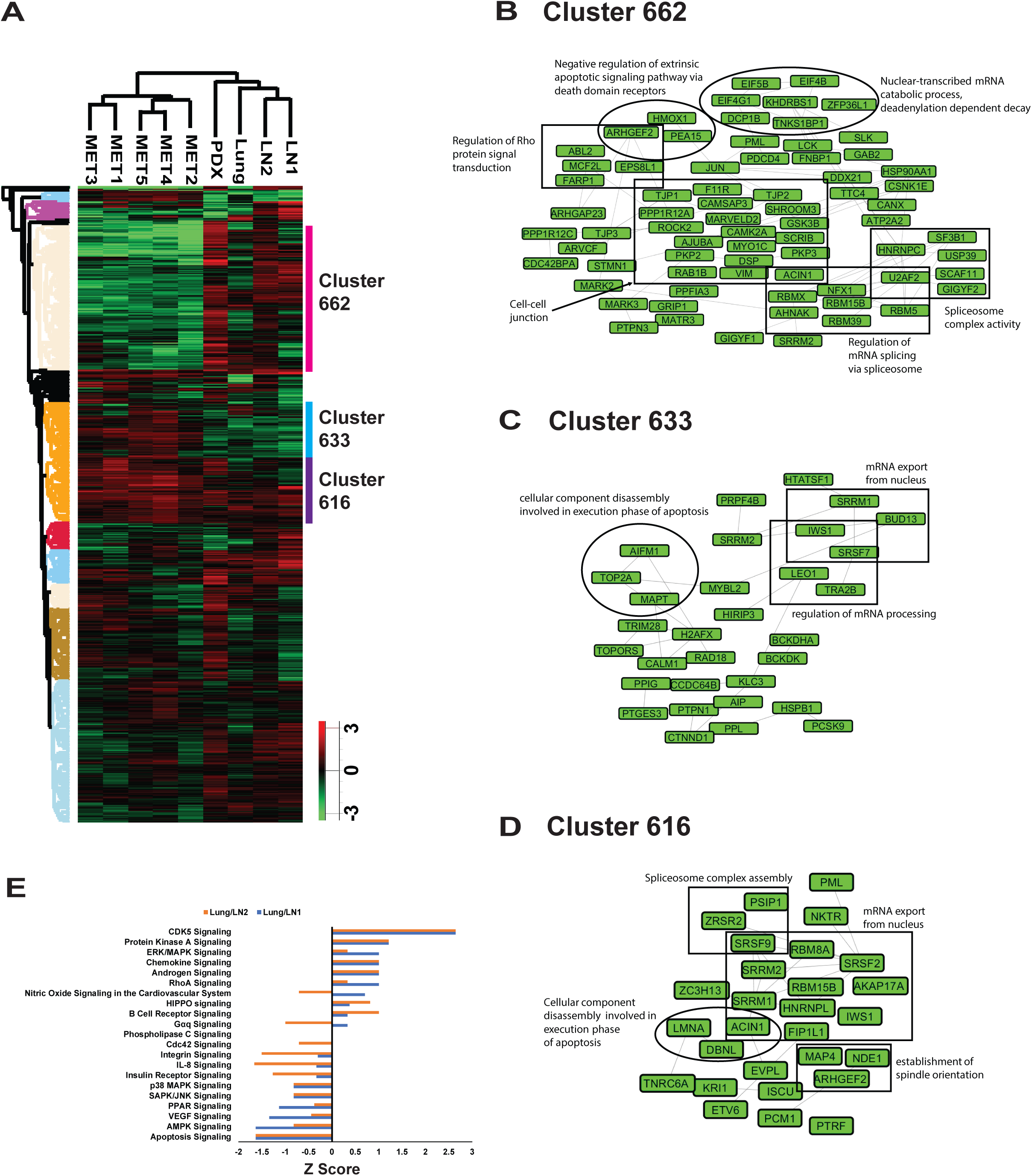
Comparison of phosphoproteome of lung and lymph node tissues. (A) Hierarchical clustering of protein phosphorylation and tissues show that autopsy samples are separated from the other tissues. Columns represent different tissue samples. Rows represent quantified phospho-sites. 3 clusters were highlighted and labelled. (B) Network of the proteins from cluster 662. (C) Network of the proteins from cluster 633. (D) Network of the proteins from cluster 616. (E) Selected top canonical pathways represented by phosphosites change from TMT experiment between lung and either lymph nodes. X-axis is the Z-score of the enriched pathways.

Ingenuity Pathway Analysis (IPA) was performed on all identified phosphorylated proteins. Among the top canonical pathways that were activated in the lung compared to lymph nodes based on the TMT phosphorylation ratio (Lung/LN2 or Lung/LN1) were CDK5, PKA, ERK/MAPK, chemokine, androgen and RhoA signalling pathways (**Fig. 5E**). Among the pathways inhibited in lung compared to the lymph nodes were apoptosis, AMPK, VEGF, PPAR, SAPK/JNK and p38/MAPK signalling pathways.

### Identification of variant peptides and mutant proteins by integrating the genomics sequencing and mass spectrometry data

We constructed patient-specific protein database based on the whole genome sequencing (5) of LN1, lung (L) and blood of this patient. We identified 78 and 23 mutant peptides from super-SILAC and TMT experiments, respectively. We validated further all the mutant peptides identified by the patient-specific database search. The mutant peptides were validated by searching through different search engines and applying a series of criteria (**Fig 6A).** In total, we validated 360 spectra corresponding to 55 germline and 6 somatic variants. Descriptions of these spectra and peptide assignments can be found in **Supplementary Table 3**. We confirmed and then annotated spectra for the somatic variants, CDK12-G879V, FASN-R1439Q and HNRNPF-A105T in the tumor lysates (**Fig. 6B, Supplementary Fig. 4, left panel)**. Heavy labelled mutant peptides were also synthesized and analyzed by mass spectrometry to match the annotated MS2 spectra for further validation (**Fig. 6B, Supplementary Fig. 4, right panel).** In addition, we searched all variant peptides for reference sequences that matched allowing for I/L isobaric substitutions but none were found.

**Figure 6.**
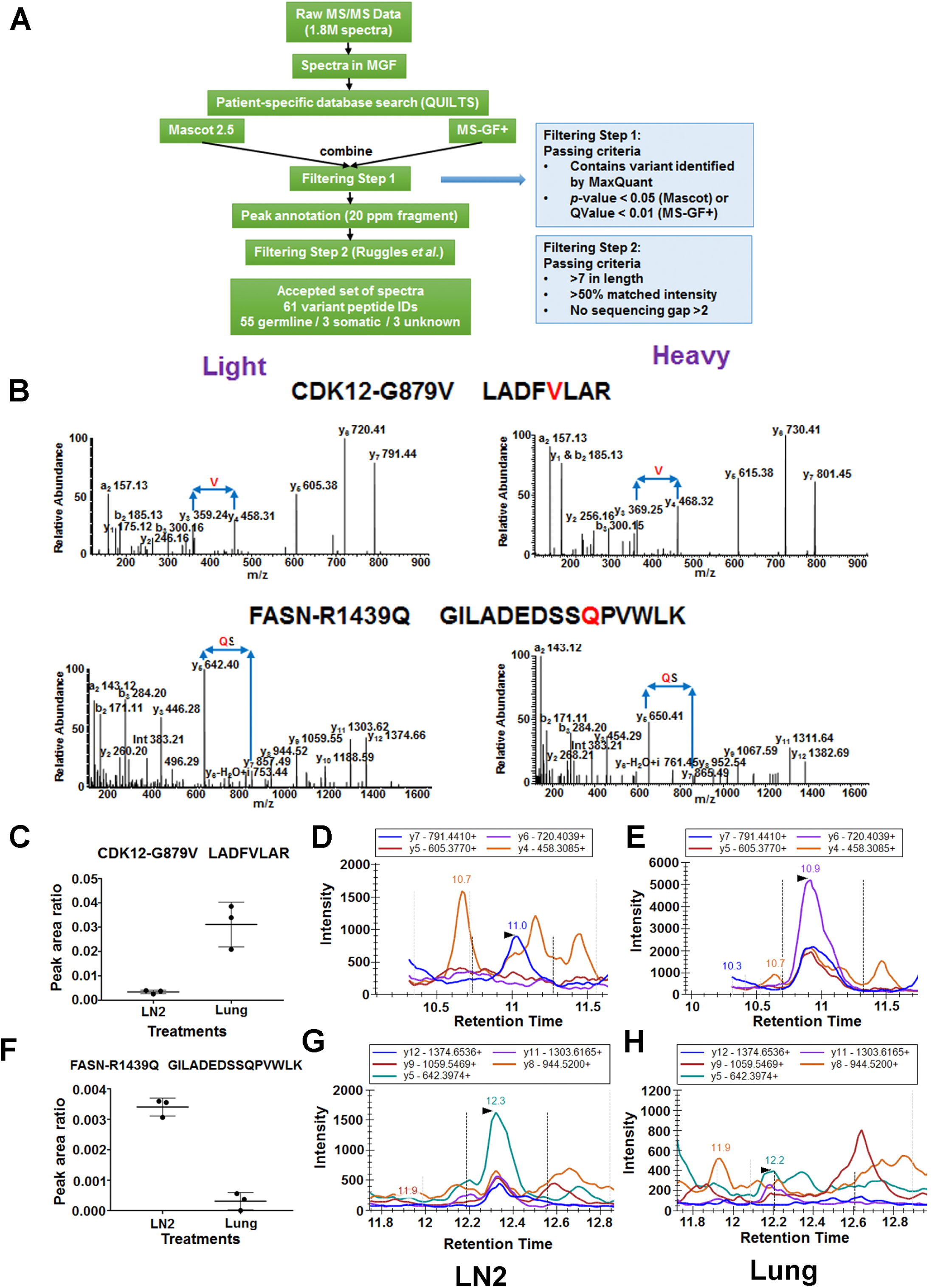
Confirmation of the mutant peptides identified from DDA data searched by the patient-specific database. (A) Flow chart of the confirmatory analysis of the MS2 spectra of all the mutant peptides. Two steps of filtering criteria were applied. (B) MS/MS spectra of two of the validated variant peptides harbouring the somatic mutations CDK12-G879V and FASN-R1439Q. On the left side are spectra of the endogenous “light” peptides identified in the lung (CDK12-G879V) and lymph node (FASN-R1439Q) lysates; on the right side are spectra of the synthetic “heavy” peptides labelled with ^13^C_6_ Arg and ^13^C_6_ Lys. (C) Relative abundance of the two mutant peptides in lung and lymph node lysates by MRM. CDK12-G879V peptide (LADFVLAR) was highly abundant in lung (L) and not detected in lymph node (LN2) while the FASN-R1439Q peptide (GILADEDSSQPVWLK) was highly abundant in LN2 and not in the lung. (D) TICs of MRM transitions of the CDK12-G879V (top) and FASN-R1439Q (bottom) mutant peptides identified in LN2 (left) and lung (L) (right) lysates. The MRM transitions of CDK12-G879V mutant peptide are identified only in the lung and those of the FASN-R1439Q mutant peptide are identified only in the lymph node.

### Targeted quantitative mass spectrometry to identify CDK12-G879V and FASN-R1439Q in corresponding metastatic lysates

We employed multiple reaction monitoring (MRM) in a triple quadrupole (QQQ) mass spectrometer to identify and relatively quantify two of the variant tryptic peptides harboring the mutations, CDK12G879V and FASN-R1439Q from the lung (L) and lymph node (LN2) metastatic sites, respectively. Two mutant peptides CDK12-G879V (LADFVLAR) and FASN-R1439Q (GILADEDSSQPVWLK) labelled with “heavy” amino acids were spiked in the tryptic peptide mix from the L and LN2 samples as internal standards for relative quantitation of their endogenous counterparts. Each sample spiked with the corresponding heavy peptide was analyzed in QQQ mass spectrometer in triplicate using the scheduled MRM method. The normalized peak area ratio of CDK12G879V peptide shows that this mutant CDK12 peptide was significantly more abundant in L compared to LN2 (**Fig. 6C)**. The TICs of the transitions from the endogenous mutated peptide can clearly be seen in L, but only at background levels in LN2 (**Fig. 6D, E**). In contrast, FASN-R1439Q peptide had higher abundance in LN2 compared to L (**Fig. 6F**) and the TICs of the transitions were only detected in LN2, but not the L (**Fig. 6 G, H**).

### Functional characterization of CDK12-G879V mutant peptide

The G879V mutation in CDK12 occurs in the DFG motif of the CDK12 kinase that is expected to destabilize the active structure of CDK12 result in a non-functional kinase. CDK12 kinase activity is required for long transcript gene expression such as of the DNA damage response (DDR) genes. Hence, we hypothesized that the non-functional G879V mutant CDK12 will increase chemotherapy sensitivity in lung adenocarcinoma cells. We knocked out the *CDK12* gene in A549 cells stably expressing Cas9 using two CRISPR sgRNAs that target exon 1 and exon 2 of *CDK12*. The CDK12 protein expression was significantly lower in both clones compared to the parental cells (**Fig. 7A**). Both knock out cell lines were more sensitive to camptothecin, a DNA topoisomerase I inhibitor that inhibits DNA replication, compared to the parental cells (**Fig. 7B**), suggesting that CDK12 function is essential for repair from DNA damage.

**Figure 7.**
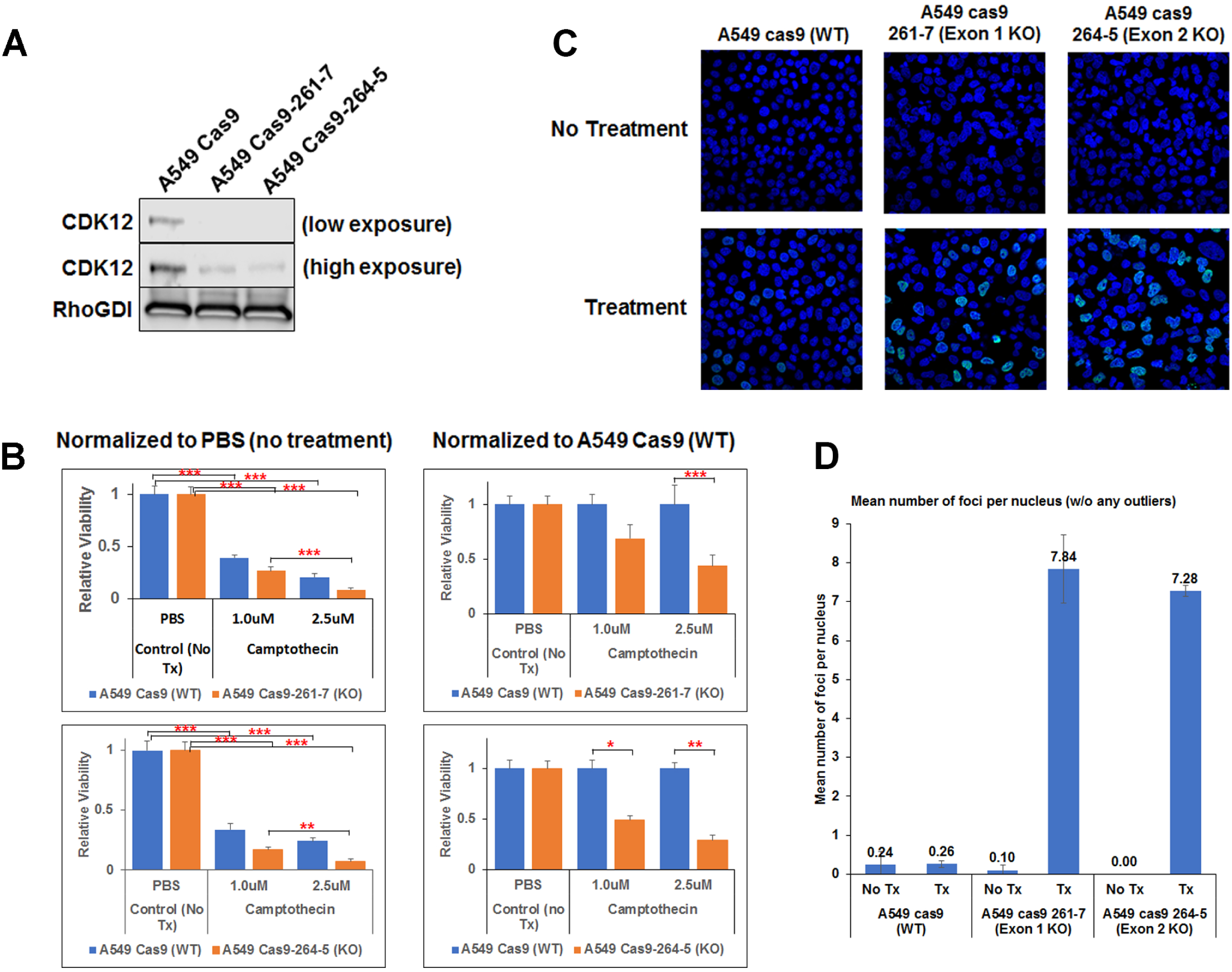
CRISPR-Cas9-mediated knockdown of CDK12 increases chemotherapy sensitivity in A549 human lung adenocarcinoma cells. (A) Western blot of CDK12 in parental A549-Cas9 control cells and stable clones expressing CRISPR gRNA targeting exon 1 (261-7) and exon 2 (264-5). There is knockdown of CDK12 in both the clones. (B) Camptothecin sensitivity of the parental and CDK12 knockout cell lines. The data is normalized to mock (PBS) treatment (left) or the with parental control cells (right). (*** p < 0.001; ** p < 0.01; * p < 0.05)

## Discussion

The overall goal of our study was to examine the heterogeneity at the level of the proteome and phosphoproteome across 11 different tumor metastases sequentially acquired by multiple biopsies and surgeries upon progression on combination HER2-directed therapy along with chemotherapy of our patient who survived with metastatic lung adenocarcinoma for over 7 years. We have previously demonstrated unprecedented mutational heterogeneity using whole genome and targeted next generation sequencing (NGS). Only <1% of somatic variants were common between the lymph node and lung metastatic sites in this patient (5). However, we demonstrated similarities in key hallmarks of carcinogenesis, such as proliferation that were manifest by different mechanisms in the lung and lymph node metastatic sites. We hypothesized that analyzing the proteome and the phosphorpoteome will provide more granular details of tumor evolution and natural history of tumor progression in this patient. To our knowledge, this is one of the initial studies to perform comprehensive genomics and mass spectrometry-based quantitative proteomics across multiple sequential biopsies of progressive lesions and an autopsy in an “exceptional responder” lung adenocarcinoma patient. Our patient responded to combination chemotherapy along with HER2-directed targeted therapy over the span of 7 years. We had demonstrated before that our patient harboured L869R ERBB2 mutation in the lymph node metastatic sites, but not the lung metastatic sites. Although ERBB2 was also amplified in both sites, the degree of amplification was far greater in the lung compared to the lymph node sites (5). Here using quantitative mass spectrometry we demonstrate that ERBB2 protein levels were 20-70 fold greater in the lung compared to the two lymph nodes. Interestingly, there was also tumor evolution and selection for higher ERBB2 protein levels in LN2 (LN2/LN1 fold change 3.2) that was procured more than 2 years after the procurement of the progressive lung metastasis by surgery and 2.5 years after procurement of the progressive lymph node LN1. The control of the multiple metastases using HER2-directed combination therapy suggests HER2, through amplification and increased protein levels in the lung and a somatic kinase domain mutation, specifically in the lymph node sites was the major driver of carcinogenesis in this patient.

One major reason why the patient might have survived such a long time with metastatic disease was that his lung metastases were essentially “cured” after the surgical removal of the progressive lung tumor, that was likely the primary lesion. At autopsy, no lung tumor metastases could be detected. One reason could be the significantly increased ERBB2 protein levels in the lung metastases and the absence of the ERBB2 L869R mutation. Hence the lung tumor metastases were more dependent on HER2 inhibition by combination targeted therapy, while the lymph node metastases were relatively resistant and hence continuously progressed. We also demonstrate the effect of the novel CDK12-G879V mutation that was present only in the lung metastasis, but not the lymph nodes in promoting chemotherapy sensitivity of the lung metastases. Here, we show convincingly that the CDK12-G879V mutant is expressed at the protein-level in the lung metastasis, but not the lymph nodes (**Fig. 6**). This mutation falls within the conserved DFG motif in the kinase domain of CDK12. A CDK12-D877N mutation in the same DFG motif has been documented in a patient with cholangiocarcinoma in TCGA and this mutation has been used as a kinase dead control in various studies (17). CDK12 in association with cyclin K has been shown to regulate transcriptional elongation of long transcript genes, such as genes involved in DNA dame response (DDR) (18, 19). CDK12 was demonstrated to be necessary for expression of long transcript genes, BRCA1, ATR, FANCI and FANCD2 upon DNA damage. This was attributed to phosphorylation of Ser-2 in the heptapeptide repeat located in the C-terminal domain (CTD) of RNA polymerase II (20). CDK12 is mutated in the kinase domain in about 3% of serous ovarian cancer patients in the TCGA. In another study depletion of CDK12 resulted in deceased BRCA1 levels, reduced RAD51 foci formation and HR repair in ovarian cancer cells (19). We hypothesized that G879V mutation within the conserved DFG motif in the kinase domain of CDK12 will result in a non-functional kinase and increase chemotherapy sensitivity of tumors. We engineered CRSPR-Cas9 mediated knockdown of CDK12 in a lung adenocarcinoma cell line, A549. Complete knockdown of CDK12 protein expression indeed increased chemotherapy sensitivity of these cells (Fig 7) that correlated with increased DNA damage foci in the knock-down cells treated with chemotherapy suggesting more DNA damage and less repair.

We compared two methodologies to perform quantitative mass spectrometry using DDA data, namely the “super-SILAC” method and the TMT labelling method. The “super-SILAC” method has been used before to quantify the proteome in human breast cancer by spiking “super-SILAC” mix of heavy labelled lysates from breast cancer cell lines (21). We have used a similar strategy, and to our knowledge, first time made a lung adenocarcinoma-specific “super-SILAC” mix of heavy labelled lysates. We developed a pooled sample of heavy labelled lysates of 12 cell lines, including 2 immortalized normal lung epithelial cell lines, 1 lung adenocarcinoma (LUAD) cell line without EGFR or KRAS mutations, 6 EGFR mutant LUAD cell lines harbouring the most common EGFR mutations, and 3 KRAS mutant LUAD cell lines. Since this pool uses some of the major subtypes of lung adenocarcinoma, this can be used to compare mass spectrometry-based proteomics experiments performed in different laboratories. The problem with “super-SILAC” data is that of missing data. Our super-SILAC strategy identified slightly more number of proteins, however, quantified proteins were similar to the TMT labelling strategy (**Fig. 1D, E**). More importantly, the correlation of the quantitation data overall was quite good between the two strategies with R^2^ values between 0.64 and 0.79 (**Fig. 2A**).

One purpose of this study was to correlate genomic and proteomic data using copy number analysis (CNA), transcriptome analysis (RNA-seq) and mass spectrometry-based quantitative proteomics. CNA clearly showed that the lung metastatic site was significantly different than 8 other lymph node metastases (**Fig. 4A**). This reinforces the unprecedented genomic tumor heterogeneity already demonstrated in this patient at the level of somatic variants, in which we had shown that there were <1% similarity of somatic variants between the lung and lymph node metastases by whole genome sequencing (WGS) (5). Adequate quality of RNA sample was available from three distinct metastases, namely LN1, L and PDX. Spearman’s correlation (range-0.28-0.52) of the RNA-seq-based FPKM values for transcript expression and either the super-SILAC or the TMT protein ratio across datasets demonstrated that there is low correlation of the transcript and protein data from the same sample and this is in the range demonstrated before in various systems.

We made patient-specific database using the WGS data published before (5) adding all germline and somatic variants identified in the lung and lymph node metastases and normal blood to the normal human RefSeq database. We used an extensive validation pipeline to confirm the identification of the variant MS/MS spectra. We also matched the spectra obtained from patient’s tumor tissue with heavy labelled variant synthetic peptides. Although, total number of germline and somatic variant peptides were around 2000, we identified only 55 germline and 6 somatic variants. Interestingly, we could identify the CDK12-G879V mutant tryptic peptide in tumor lysates from the lung metastasis without any enrichment. This could be a result of overexpression of the mutant protein due to documented amplification of the *ERBB*2 and *CDK12* locus. The overall low percentage of variant peptide detection by mass spectrometry has been documented before (6-9). This is likely a result of limited peptide coverage of the individual proteins by current mass spectrometry technologies in global proteomics experiments. In spite of this limited detection of variant peptides, any variant peptide that is detected by mass spectrometry is likely to be important since the variant is expressed at the protein level. Here, we also developed an MRM assay for the CDK12G879V novel variant to perform quantitative estimation of expression of the variant peptide/protein. Such assays against variant peptides would allow quantitation of mutant proteins from limited amount of biopsy samples.

Proteomic and phosphoproteomic heterogeneity has been demonstrated by quantitative mass spectrometry in pancreatic cancer from metastatic sites from a single patient procured at autopsy. Interestingly, an AXL inhibitor was more effective in lung and liver metastatic sites compared to the one from the peritoneum, suggesting that there was heterogeneity in activation of signalling pathways in different metastatic sites (22). Our study involves multiple metastatic sites procured during the treatment course over 7 years, including at autopsy. We also demonstrate the importance of integrated proteogenomic analyses in such studies to identify variant peptides in mass spectrometry data. We demonstrate how CDK12G879V mutant specific to the lung metastatic site likely contributed to the superior sensitivity of the lung metastatic sites to combination of chemotherapy along with HER2-targeted therapy. We also developed an MRM assay to quantify the CDK12G879V mutant peptide. We could identify this variant peptide in the lung lysates without any enrichment, likely due to high mutant protein expression and confirming the amplification of *CDK12* from whole genome sequencing and CNV-seq data.

In summary, here, we have shown, the feasibility of multiple biopsies through a patient’s treatment course culminating in an autopsy upon death of an “exceptional responder” lung adenocarcinoma patient. The global proteomics and phosphoproteomics performed here along with CNA and transcript expression described in this study and the NGS analyses published previously illuminate the unique tumor evolution of this patient. Most importantly, our study highlights the functional significance of one of the most relevant somatic variants, the CDK12-G879V mutation that could be identified directly at the peptide-level by both DDA and MRM mass spectrometry. This variant may have influenced the overall prognosis and treatment response in this patient.

## Acknowledgements

This work was supported by NCI Intramural Program support to U.G and also partly from the CPTAC grant U24 CA210972.

## Data submission

The MS proteomics data in this study has been deposited in ProteomeXchange Consortium (http:proteomecentral.proteomeexchange.org) via the PRIDE partner repository with the dataset identifier PXD010779. The reviewer account details are as follows: username: password: mDAxieid.

---

**Supplementary Figure 1.**
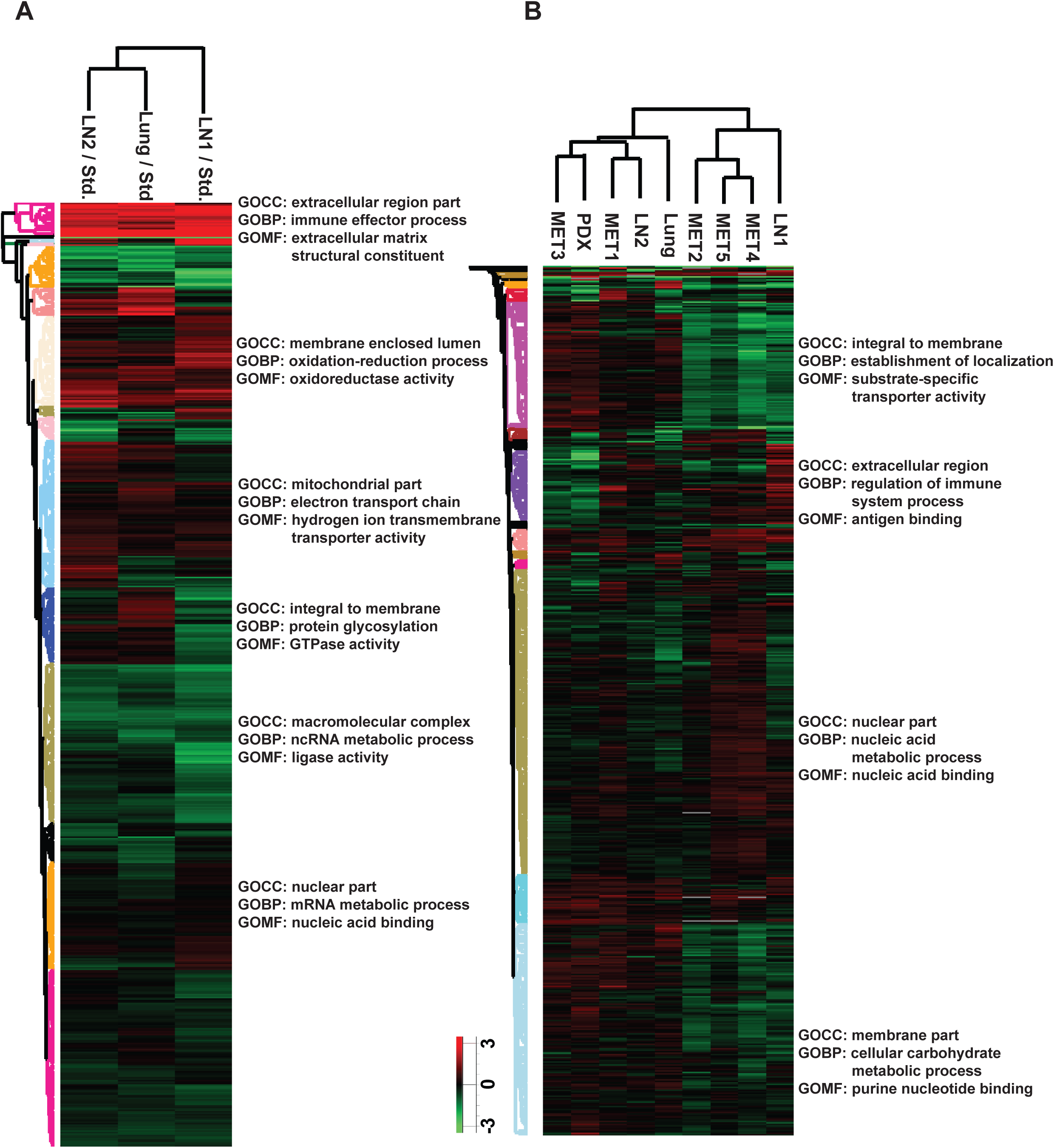
Hierarchical clustering of patient tumor tissue proteome. (A) Hierarchical clustering of proteins and three biopsy tumor tissues from super-SILAC experiment shows that LN1 is separated from lung and LN2 tissues. (B) Hierarchical clustering of proteins and nine tissues from TMT experiment. Two clusters of tissues are grouped based on the protein expression.

**Supplementary Figure 2.**
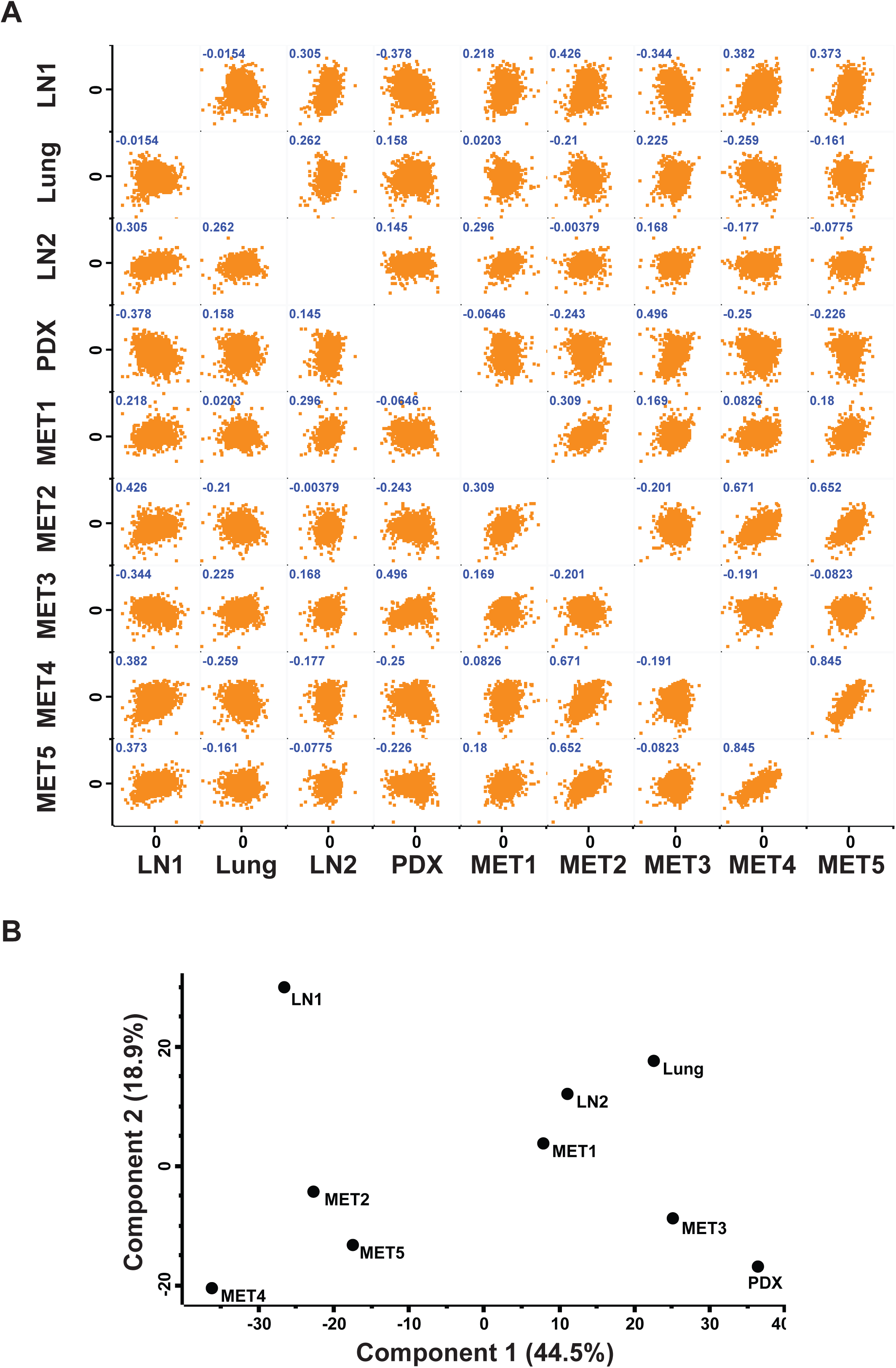
Comparison of lung and lymph node tissue proteome. (A) Correlation of protein expression between the three tumor biopsies, one PDX, and five autopsy samples from the TMT experiment. (B) Principal component analysis of the tissues based on the protein expression shows two clusters, the same as the hierarchical clusters.

**Supplementary Figure 3.**
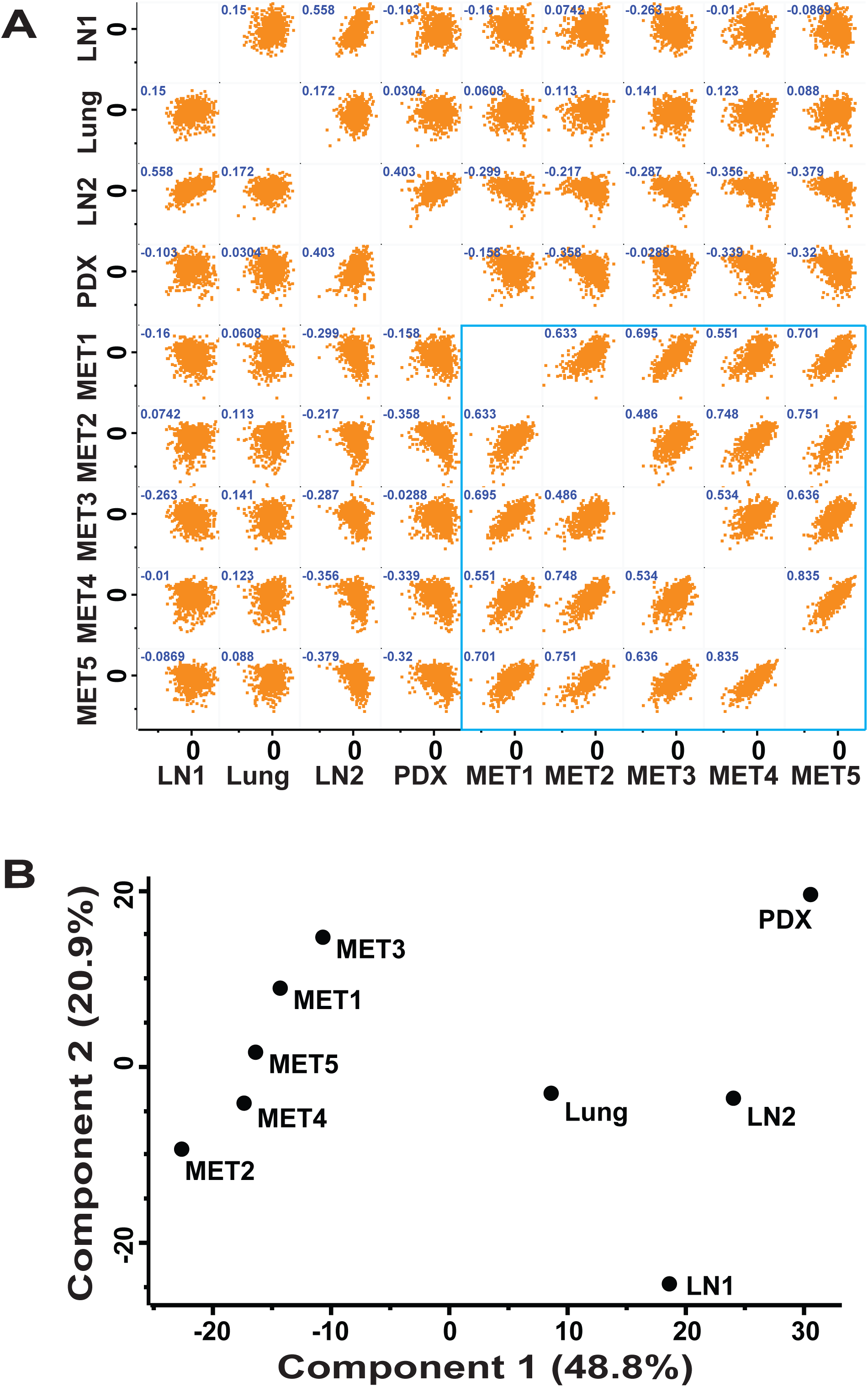
Comparison of phosphoproteome of lung and lymph node tissues. (A) Correlation of protein phosphorylation between the three tumor biopsies, one PDX, and five autopsy samples from the TMT experiment. The five autopsy samples show higher correlation than the other samples (the blue box). (B) Principal component analysis of the tissues based on the protein phosphorylation differentiates the autopsy tumor tissues from the other tumor biopsies and PDX sample.

**Supplementary Figure 4.**
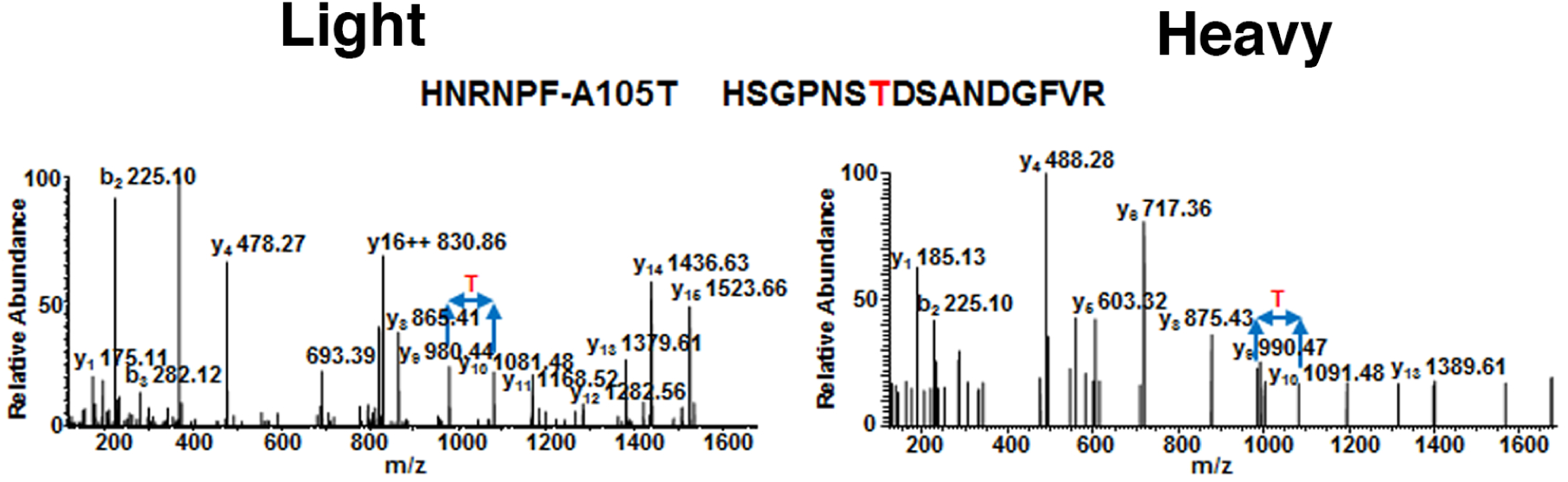
Confirmation of the mutant peptide, HNRNPF-A105T identified from DDA data searched by the patient-specific database. MS/MS spectrum of the light endogenous mutant peptide (left panel) and the heavy amino acid labelled synthesized peptide (right panel) show the presence of the mutation in HNRNPF.

